# iDRKAN: Interpretable miRNA-Disease Association Prediction Based on Dual-Graph Representation Learning and Kolmogorov–Arnold Network

**DOI:** 10.1101/2025.02.11.637780

**Authors:** Yangfeng Zhu, Yongxian Fan, Guicong Sun, Xueping Li

## Abstract

Accurately identifying miRNA-disease association (MDA) is of great importance in biomedical research and clinical applications. However, most existing computational methods rely on similarity, making it difficult to effectively capture the deep semantic information among heterogeneous nodes in complex networks. In addition, the inherent “black-box” nature of traditional deep learning models leads to a lack of transparency in their decision-making process. Therefore, we propose an interpretable MDA prediction method (iDRKAN) based on dual-graph representation learning and Kolmogorov-Arnold Network. First, iDRKAN constructs similarity views and meta-path views based on the similarity matrix and association matrix respectively, and learns higher-order feature representation of each view by graph convolutional network (GCN). Subsequently, the multi-channel attention (MCA) mechanism is introduced to adaptively fuse the contextual information of each similarity view in different convolutional layers, and the semantic layer attention (SLA) mechanism is used to integrate the semantic information contained in different meta-path views. Next, the contrastive learning strategy is used to optimize the consistency between the dual-graph representation. Finally, the dual-graph features are weighted fused, and fed into the interpretable Kolmogorov-Arnold Network (KAN) for prediction. Experimental results on two public datasets show that iDRKAN significantly outperforms existing computational approaches in multiple performance indicators. In addition, experiments with different classifiers verify that iDRKAN achieves a good balance between prediction performance and interpretability. The case study further demonstrates the effectiveness of iDRKAN in mining potential associations.

## Introduction

MicroRNA (miRNA) is a subset of non-coding RNA, usually an endogenous initiating short RNA molecule about 22 nt in length [1]. miRNA is not only involved in a variety of biological processes such as cellular proliferation and differentiation [2], but also associated with many human diseases, such as cancer [3, 4, 5], cardiovascular disease [6], and other complex diseases. For example, compared with normal tissues, many miRNA transcription is upregulated in papillary thyroid cancer tumors. Among them, miR-221, miR-222, and miR-146 are the most significantly upregulated, and they can effectively distinguish papillary thyroid cancer from normal thyroid [7]. Therefore, accurate identification of disease-related miRNA facilitates the development of disease research from single-factor analysis to systems biology and precision medicine.

To date, most MDAs are derived from biological wet experiments, which are both time-consuming and costly [8]. With the development of intelligent computing, using computational methods to mine potential MDAs is both time-saving and labor-saving. Generally, these computational methods can be broadly categorized into two groups: similarity-based methods and machine learning-based methods. Similarity-based methods prioritize disease-related miRNAs, assuming that functionally similar miRNAs are associated with phenotypically similar diseases [9, 10]. Typical similarity calculation methods in MDA prediction include disease semantic similarity [11], miRNA functional similarity [12], and Gaussian interaction profile (GIP) kernel similarity [13]. For example, Han et.al [14] constructed multiple similarity matrices and fused the prediction results of each similarity matrix based on graph semi-supervised learning. Wang et.al [15] proposed a nonlinear fusion strategy to mitigate the noise caused by multi-source data and its impact on network quality. Wang et.al [16] introduced genes to construct a multilayered heterogeneous network and predicted potential MDA through genes with similar functions. Li et.al [17] used the similarity matrix and known association matrix to achieve matrix completion. Feng et.al [18] completed matrix reconstruction by aggregating first-order neighbor information based on node reliability. Chen et.al [19] combined domain constraint and matrix completion as similarity-assisted information for MDA prediction. Although similarity-based methods can effectively utilize known association information, their performance depends on the choice of similarity metric and the quality of feature representation.

With the development of machine learning technology, its application in bioinformatics has become more and more widespread. For instance, Liu et.al [20] first used a stacked autoencoder to extract potential feature representation and subsequently combined them with XGBoost to achieve prediction. In addition, extreme imbalance of data can cause the prediction model to be biased towards the majority class, thus affecting the prediction accuracy. To this end, Dai et.al [21] balanced the data through a resampling technique and combined the soft voting strategy of ensemble learning for prediction. Wang et.al [22] first trained the autoencoder in an unsupervised manner and then used balanced samples for supervised optimization. Liu et.al [23] generated different low-dimensional embeddings through dual autoencoders and used the random forest to implement the classification prediction. In recent years, with the increasing demand for complex biological network analysis, graph-based methods have shown excellent performance in processing high-dimensional sparse data. Graph Convolutional Neural Networks (GCNs) [24] are particularly suitable for processing non-Euclidean data structures. They can efficiently capture complex relationships between neighbors of different orders and automatically learn the embedding representation of nodes. For example, Peng et.al [25] extracted consistent representation among different hypergraphs by hypergraph convolution and contrastive learning strategy. Ning et.al [26] introduced virtual nodes to construct a heterogeneous hypergraph to enrich link information. Lou et.al [27] constructed a multimodal network based on the similarity matrix and association matrix, and aggregated multi-order neighbor information in the network through GCN. Tang et.al [28] extracted the embedding representation of multiple views via GCN, and then used the multi-channel attention mechanism to fuse the feature matrices of each view. He et.al [29] constructed miRNA-disease and disease-gene sub-networks by gene to alleviate the problem of data sparsity. Although GCN’s graph information aggregation capability is efficient, it is susceptible to over-smoothing and data noise. Over-smoothing reduces the discriminability of node representations, while data noise may mislead model learning.

Most current approaches rely on the similarity of homogeneous nodes (such as disease semantic similarity), while often ignoring the deep semantic information between heterogeneous nodes. Moreover, the “black-box” nature of these models reduces the transparency of the decision-making process. To address these limitations, we propose an interpretable MDA prediction method (iDRKAN) based on dual-graph representation learning and Kolmogorov-Arnold Network. Specifically, iDRKAN first constructs similarity views and meta-path views based on similarity metrics and known association, and extracts the corresponding graph embedding via GCN. Subsequently, MCA and SLA are used to fuse the features from each similarity view and different meta-path views. Next, contrastive learning is further applied to optimize the consistent representation of dual-graph. Finally, the weighted fused dual-graph features are input into KAN for prediction. We evaluate the effectiveness of iDRKAN on two public datasets. To further validate the balance between prediction performance and interpretability of iDRKAN, we also conducted experiments using the iDRKAN framework in combination with various classifiers. The overall architecture of iDRKAN is illustrated in Fig. 1.

**Fig. 1.**
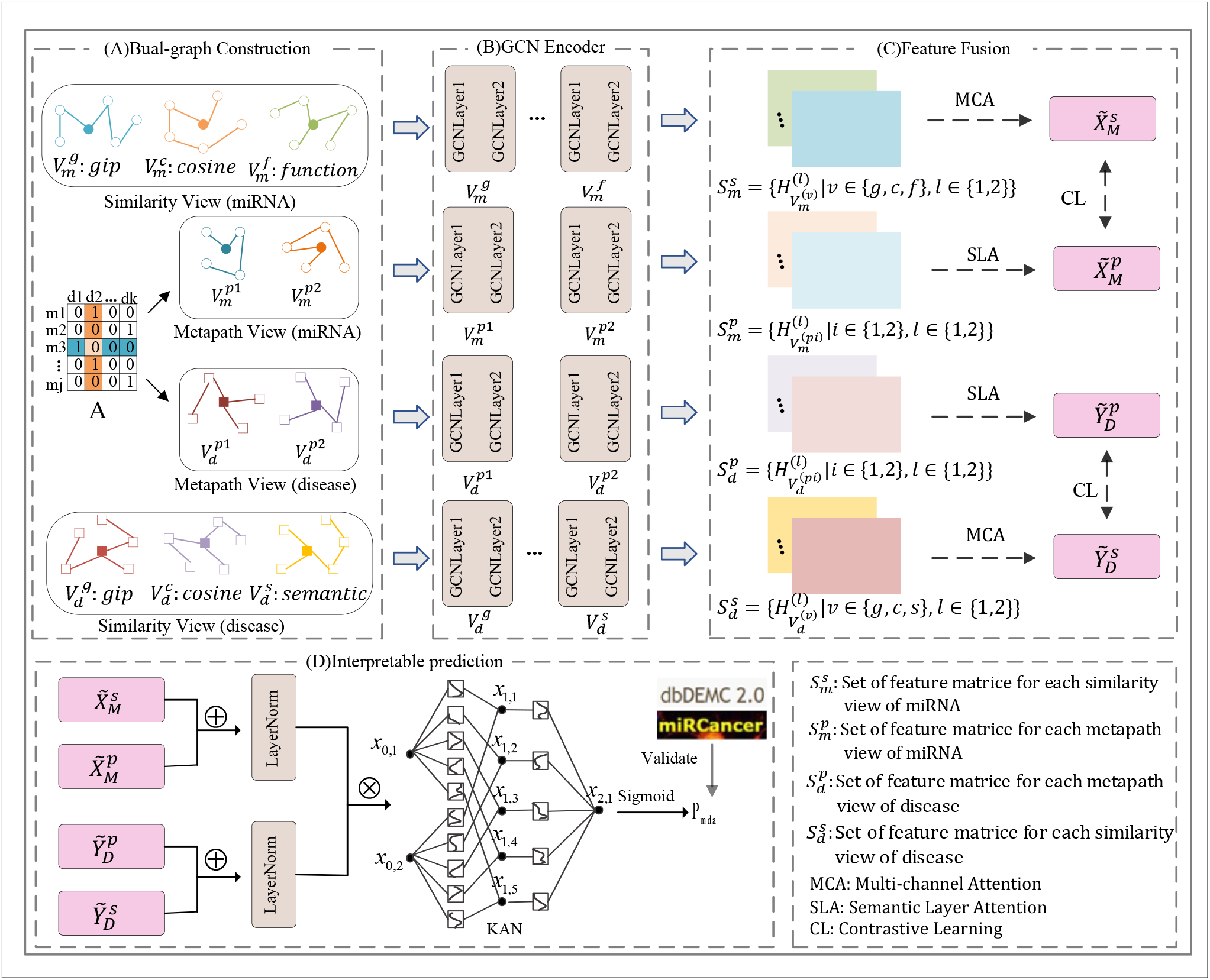
The overall framework of iDRKAN. (A) Construction of similarity and meta-path view. (B) Extraction of high-order representation from each similarity and meta-path vies using GCN. (C) MCA is used to fuse the contextual information of each similarity view, and SLA is utilized to integrate the semantic information contained in different meta-path views. (D) Kolmogorov–Arnold Network is used for interpretable MDA prediction.

## Materials and Methods

### 2.1 Related Materials

HMDD v4.0 [30] and miR2Disease [31] were used as the baseline datasets in this study. HMDD v4.0 contains 1817 miRNAs, 2360 diseases, and 53,530 experimentally validated MDA records. miR2Disease contains 349 miRNAs, 163 diseases, and 3,273 known associations. miRTarBase [32] focuses on collecting experimentally validated miRNA target genes, ensuring that the miRNAs used are experimentally confirmed. The Disease Ontology (DO) database [33] provides a comprehensive and standardized classification of diseases, with each disease corresponding to a unique Disease Ontology ID (DOID). In addition, we downloaded the Medical Subject Headings (MeSH) [34] or standardizing disease names. To ensure consistency across different data sources, we excluded diseases that were not categorized in MeSH class C or lacked a DOID identifier. Also, we removed miRNAs that were not included in the miRTarBase database. Finally, the HMDD v4.0 dataset contains 728 miRNAs, 884 diseases, and 13,511 MDA records. The miR2Disese dataset involves 1,376 associations for 215 miRNAs and 110 diseases. We define the MDA matrix as 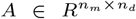, where *n*_*m*_ and *n*_*d*_ represent the number of miRNA and disease respectively.

### 2.2 Dual-Graph Construction

The similarity view of miRNA in this study includes GIP kernel similarity 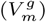, cosine similarity 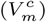, and functional similarity 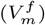. The similarity view of disease includes GIP kernel similarity 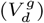, cosine similarity 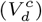, and functional similarity 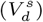. Please refer to the Supplementary file for the detailed calculation process.

Traditional similarity views are difficult to effectively capture the deep semantic association contained in a complex network. For this reason, we construct the meta-path view that aims to fully extract the deep semantic information in the network through multi-hop paths [35]. Specifically, a meta-path is denoted as 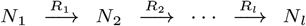, and the composite relationship between nodes *N*_1_ and *N*_*l*_ is represented as *R* = *R*_1_ ◦ *R*_2_ ◦ · · · ◦ *R*_*l*_, where ◦ is the relational conformity operator. In this paper, the meta-path of miRNA contains miRNA-disease-miRNA (MDM) and miRNA-disease-miRNA-disease-miRNA (MDMDM). The meta-path of disease includes disease-miRNA-disease (DMD) and disease-miRNA-disease-miRNA-disease (DMDMD). The meta-paths, 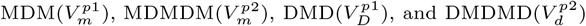, are calculated as follows:

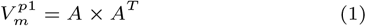

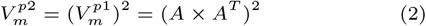

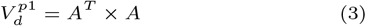

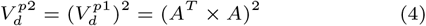

Fig. 2 shows an example of meta-path 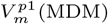 generation. In the meta-path matrix calculated in Fig. 2(C), the element 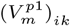 represents the number of paths between miRNA *m*_*i*_ and *m*_*k*_ connected by the meta-path MDM. For example, 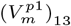 indicates that there is an MDM path between miRNA *m*_1_ and *m*_3_, which means that *m*_1_ and *m*_3_ are connected through the commonly associated disease *d*_2_ are connected through the commonly associated disease (which can be verified in both Fig. 2(A) and Fig. 2(B)). The diagonal element 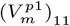 indicates that miRNA *m*_3_ has two different MDM paths returning to itself.

**Fig. 2.**
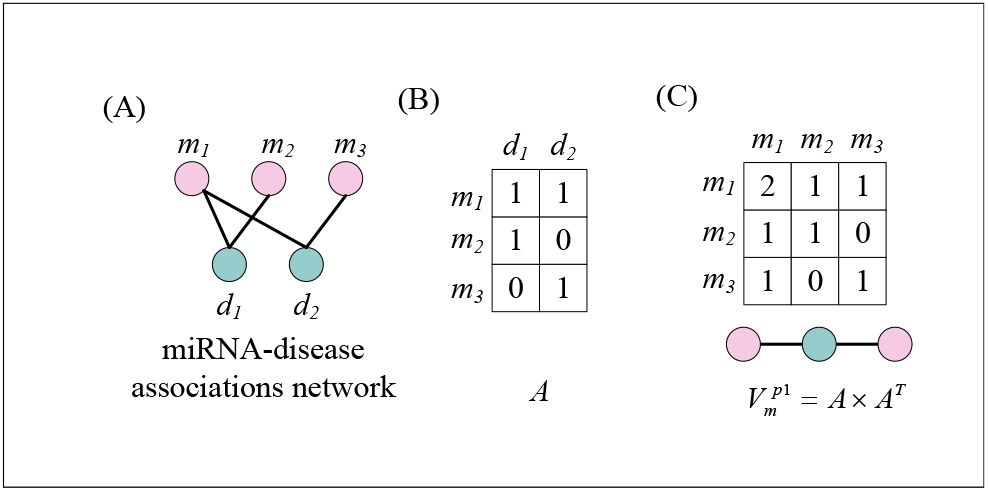
Example of the generation of the meta-path 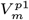.

### 2.3 Dual-Graph Representation Learning)

GCN has shown excellent performance in complex network node representation learning by its superior feature aggregation capability. In this section, we will introduce how to utilize GCN to efficiently learn node representations in similarity view and meta-path view to fully explore the contextual information and deep semantic associations in the network. Taking miRNA as an example, we define the single-layer GCN operation according to the information aggregation and update mechanism of GCN as follows:

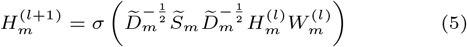

where *σ*(·) is the ReLU activation function, 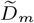 is the degree matrix of 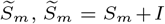 and *I* is the identity matrix. 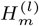 is the feature matrix of the L-th layer. 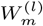 denotes the learnable weight matrix. For the similarity view 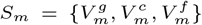, and the meta-path view 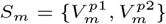.

#### 2.3.1 Extraction and Fusion of Similarity View Feature

The feature representation of the miRNA similarity view learned in each convolutional layer is as follows:

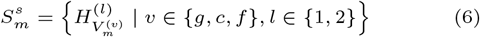

where 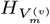 is the feature matrix of view 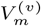 at the l-th layer.

Different similarity views may contain different contextual information, so we introduce the Multi-Channel Attention (MCA) mechanism [36] to fuse the features learned from different similarity views at each convolutional layer. Specifically, MCA stacks all feature matrices into a tensor and considers the tensor as an image. Each feature matrix can be regarded as a channel on the image. First, we generate a channel description vector based on global average pooling:

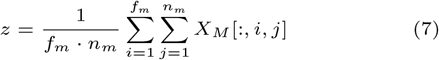

where *f*_*m*_ is the feature dimension, 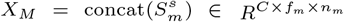, and *C* is the number of channels.

Then, a multi-layer perceptron (MLP) is used to perform the nonlinear operation on z to learn the channel attention weight:

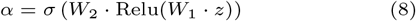

where *W*_1_ and *W*_2_ are the learnable parameters and *σ* is the sigmoid activation function. Finally, *α* is applied to each channel of *X*_*M*_ to achieve weighted fusion:

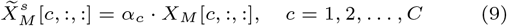

where 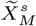 is the result of feature fusion from each similarity view of miRNA. Similarly, the feature fusion result of multiple similarity views of disease can be represented as 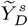.

#### 2.3.2 Extraction and Fusion of Meta-path View Feature

The feature representation of the miRNA meta-path view learned in each convolutional layer is as follows:

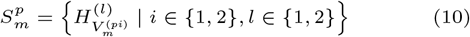

where 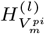 is the feature matrix of view 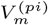 at the l-th layer.

To fuse the semantic information contained in different meta-paths, inspired by Tian et.al [37], we use Semantic Layer Attention (SLA) to integrate the features learned from different meta-path views at each convolutional layer. The feature fusion result 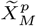 of each meta-path view of miRNA is calculated as follows:

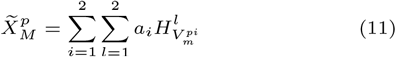

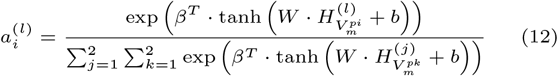

where *W* and *b* are learnable parameters, and a denotes the attention weight vector. Similarly, the feature fusion result of multiple meta-path views of disease can be expressed as 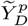

### 2.4 Contrastive Learning Strategy

For similarity fusion feature 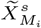 and meta-path fusion feature 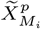 of miRNA *m*_*i*_, we first project them to the same latent space through a linear layer. Subsequently, we employ the contrastive learning strategy to enhance the consistency of cross-view representation to mitigate the heterogeneity of feature distribution in different views.

Traditional methods usually take the same miRNAs in different views as positive sample pairs when screening positive sample pairs, but this strategy may be difficult to capture more complex semantic relationships. In this study, we define that if there is a certain number of meta-paths between two different miRNAs (the threshold of the number of meta-paths in this study is set to 10), they can be regarded as positive sample pairs. Assuming that all positive and negative sample pairs are denoted as 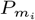 and 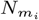 respectively, the contrastive loss of the miRNA similarity view is calculated as follows:

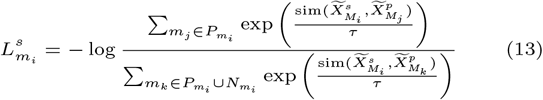

where sim(·) is is the cosine similarity between miRNA *m*_*i*_ and *m*_*j*_, and *τ* is the temperature coefficient (*τ* is set 0.7 in this study).

Similarly, the contrastive loss for the miRNA meta-path view is defined as follows:

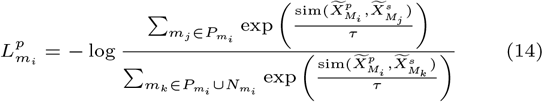

The total contrastive loss for miRNA learning representation can be formulated as follows:

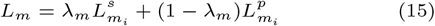

Similarly, the total contrastive loss for disease learning representation can be defined as follows:

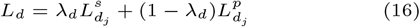

where *λ*_*m*_ and *λ*_*d*_ are the contribution coefficients of balancing the similarity view and the meta-path view (*λ*_*m*_ and *λ*_*d*_ are both set to 0.5 in this study).

### 2.5 Kolmogorov-Arnold Networks(KANs)

Inspired by the above theory [38], KAN [39] solves the constraints of the Kolmogorov-Arnold representation theorem by stacking KAN layers containing an arbitrary number of nodes. KAN allows for arbitrary widths and depths, and the original theory can be regarded as a special case of KAN with a middle layer having two layers and 2*n*+1 nodes. A single-layer KAN is defined as follows:

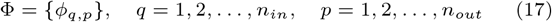

where *ϕ*_*q,p*_ is a univariate trainable activation function, and *n*_*in*_ and *n*_*out*_ represent its input and output respectively. Given a *n*_*in*_ − dimensional input vector 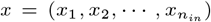, the output layer of KAN is formulated as follows:

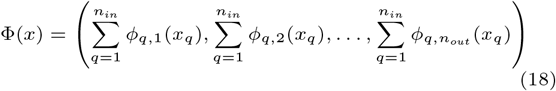

The general L-layer KAN can be then defined as follows:

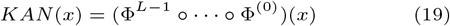

where 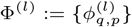 is the l-th layer KAN.

### 2.6 Prediction and Explainability Analysis

In recent years, neural networks have significantly improved in performance compared to traditional machine learning algorithms, but their intrinsic complexity leads to a lack of interpretability in the model. Therefore, in this study, KAN with display function expression structure is used as the classifier, aiming to increase the decision transparency of the model according to the learnable symbolic activation function within KAN. We combine the optimized features from the similarity view and the meta-path view, and then input them into KAN to compute the corresponding association probabilities:

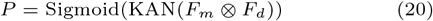

where ⊗ is the Hadamard product. 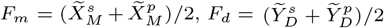.

In the training phase, we use binary cross entropy as the loss function to calculate the prediction loss *L*_*p*_. Combining the two loss functions in contrastive learning (Eq. 15 and Eq. 16), the total loss is defined as follows:

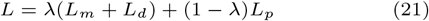

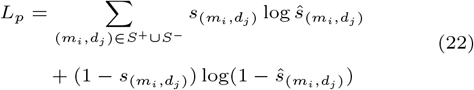

where *λ* is the contribution coefficient to measure the contrastive loss and prediction loss (*λ* is set to 0.5 in this study). *m*_*i*_ and *d*_*j*_ are miRNA and disease samples, and *S*^+^ and *S*^−^ are the positive and negative sample subsets for training. If (*m*_*i*_, *m*_*j*_) ∈ *S*^+^ then 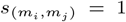, otherwise 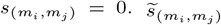 is the prediction probability.

The overall process of KAN is displayed in Fig. 3. In the initialization stage (A), the B-spline functions are randomly initialized. Subsequently, in the training phase (B), these B-spline functions gradually fit the nonlinear relationship between the input features (f1-f4) and the output. In addition, the pre-pruning stage (C) will remove the B-spline functions that contribute less to the output, thereby simplifying the network structure. To alleviate the possible performance degradation caused by the pruning operation, the fine-tuning phase (D) further optimizes the model. Finally, in the symbolization stage (E-H), the B-spline functions are replaced with mathematical expressions (such as trigonometric functions, exponential functions, etc.), and this symbolization technique significantly improves the decision transparency of the model. Specifically, assume that the two input features of miRNA-disease embedding pairs are *f*_1_ and *f*_2_, and the goal is to predict the association probability *y*. In traditional neural networks, the feature relationships are usually hidden in complex weight matrices and nonlinear activation functions, which are difficult to interpret directly. In contrast, KAN can express the relationship between the input features through a learnable symbolic activation function:

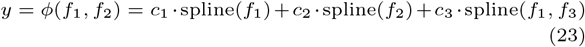

where spline(*f*) is the learnable B-spline function to capture the nonlinear effect of *f*. *c*_1_, *c*_2_, and *c*_3_ are the learnable weight. spline(*f*_1_, *f*_2_) denotes the interaction effect between features *f*_1_ and *f*_2_. Eq. 24 further illustrates the substitution of the B-spline function with a specific mathematical expression:

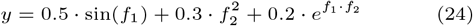

**Fig. 3.**
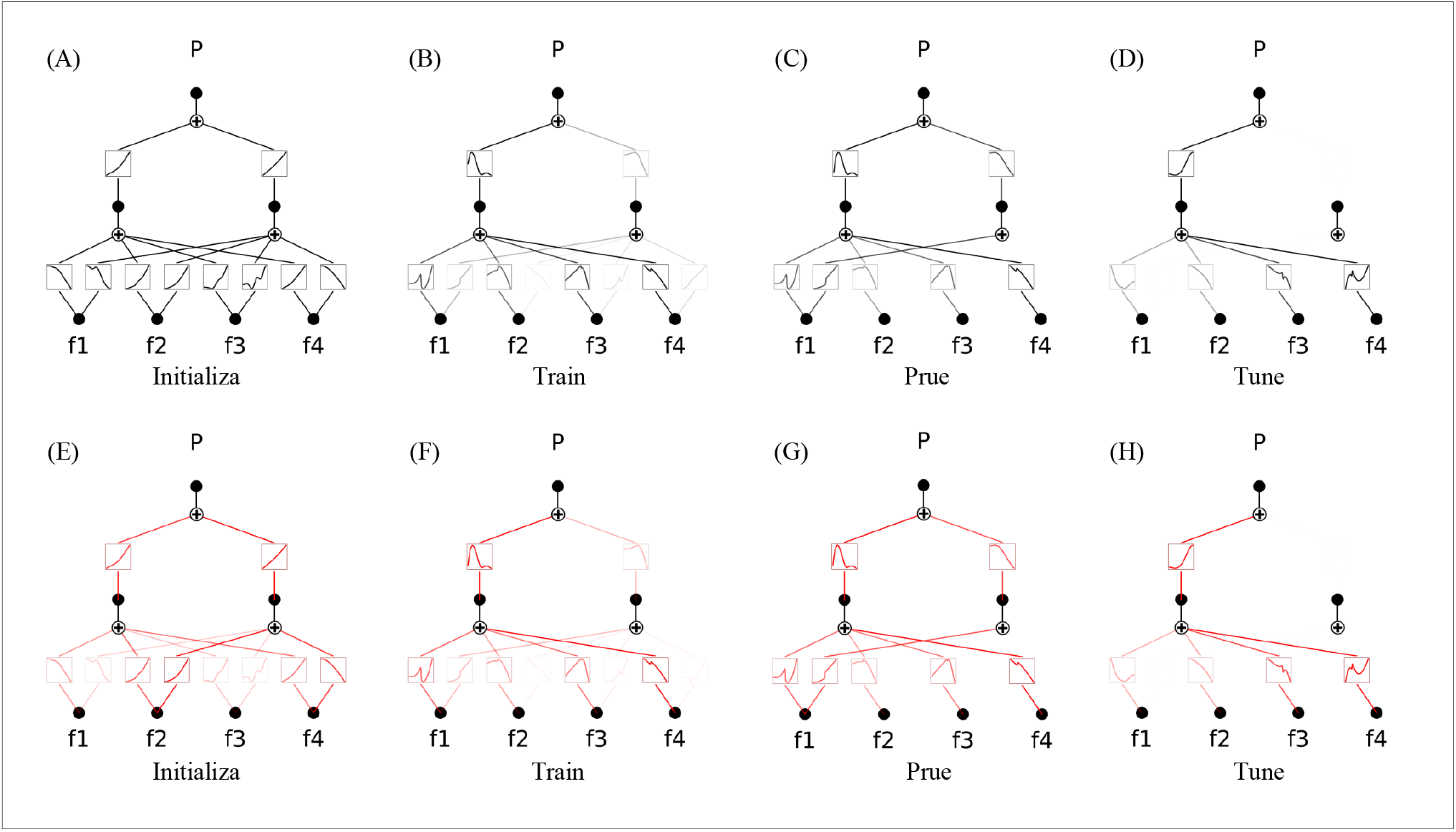
The overall process of KAN before (top row) and after (bottom row) symbolization. The black activation function is the B-spline function, while the red activation function indicates the symbolization function after replacement. Edge brightness is proportional to the effect strength.

This mathematical expression makes the contribution of each feature to the prediction result more transparent by specifying the relationship between the features, which facilitates the understanding of the decision-making process of the model.

## Experiment and Result

### 3.1 Experiment Setup

We evaluate the performance of iDRKAN using ten-fold cross-validation. Specifically, all known associations are treated as positive samples and randomly divided into ten subsets. Nine of these subsets are selected as the training set each time, and the remaining subset is used as the test set. This process is repeated ten times to ensure that each subset is used as an independent test set once. The experimental parameters are set as follows: learning rate (0.001), weight decay (1e-5), number of convolutional layers (2), convolutional dimension (64), number of KAN layers (3), and three-layer KAN dimensions (64 → 32, 32 → 16 and 16 → 1). Performance evaluation metrics include AUC, AUPR, ACCURACY, MCC, and F1-SCORE.

### 3.2 Performance Evaluation

#### 3.2.1 Comparison with Baseline Methods

To demonstrate the superiority of the proposed method, we compare it with four baseline methods, including AMHMDA based on heterogeneous hypergraphs [26], MHCLMDA based on multi-hypergraph contrastive learning [25], MINIMDA based on hybrid domain information [27], and MMGCN based on multi-view multi-channel attention fusion [28]. To ensure the fairness of the experiments, all the above models are implemented under the same experimental conditions, including the same similarity metrics and dataset partitioning methods.

As shown in Table 1, although the F1-SCORE of iDRKAN on the dataset miR2Disease is slightly lower than AMHMDA, it outperforms the other baseline methods on all other assessment metrics. Table 2 presents the performance comparison of iDRKAN with other methods on the HMDD v4.0 dataset. iDRKAN achieves an AUC of 0.9598 on this dataset, demonstrating a significant performance improvement compared to the evaluation results on the miR2Disease dataset. The excellent performance of iDRKAN can be attributed to three key factors. First, by constructing the meta-path view containing deep semantic information and optimizing the consistency of representations in different views leveraging contrastive learning, the heterogeneity of feature distribution across views is effectively reduced. Second, MCA effectively fuses the contextual information of different similarity views, while SLA fully integrates the semantic information contained in each meta-path view, enabling more comprehensive feature representations. Finally, KAN, with its learnable symbolic activation function, can flexibly approximate the complex nonlinear relationship between the input features and the target, thereby enhancing the prediction accuracy of the model. Fig. 4(A-B) and Fig. 4(C-D) present the ROC and PR curves of iDRKAN and its baseline method on the HMDD v4.0 and miR2Disease datasets respectively.

**Table 1.** Comparison of iDRKAN and baseline methods on the miR2Disease dataset.

| Methods | ACCURACY | MCC | F1-SCORE | AUROC | AUPR |
| --- | --- | --- | --- | --- | --- |
| AMHMDA | 0.8370 | 0.6761 | **0.8432** | 0.8904 | 0.8812 |
| MHCLMDA | 0.8297 | 0.6599 | 0.8266 | 0.8704 | 0.9038 |
| MINIMDA | 0.7774 | 0.5642 | 0.7960 | 0.8632 | 0.8584 |
| MMGCN | 0.7971 | 0.6181 | 0.8217 | 0.8824 | 0.8748 |
| **Our** | **0.8406** | **0.6812** | 0.8417 | **0.9063** | **0.9095** |

**Table 2.** Comparison of iDRKAN and baseline methods on the HMDD v4.0 dataset.

| Methods | ACCURACY | MCC | F1-SCORE | AUROC | AUPR |
| --- | --- | --- | --- | --- | --- |
| AMHMDA | 0.8728 | 0.7463 | 0.8755 | 0.9343 | 0.9244 |
| MHCLMDA | 0.8868 | 0.7742 | 0.8846 | 0.9352 | 0.9468 |
| MINIMDA | 0.8938 | 0.7887 | 0.8965 | 0.9447 | 0.9466 |
| MMGCN | 0.8661 | 0.7358 | 0.8724 | 0.9224 | 0.9062 |
| **Our** | **0.9038** | **0.8078** | **0.9050** | **0.9598** | **0.9591** |

**Fig. 4.**
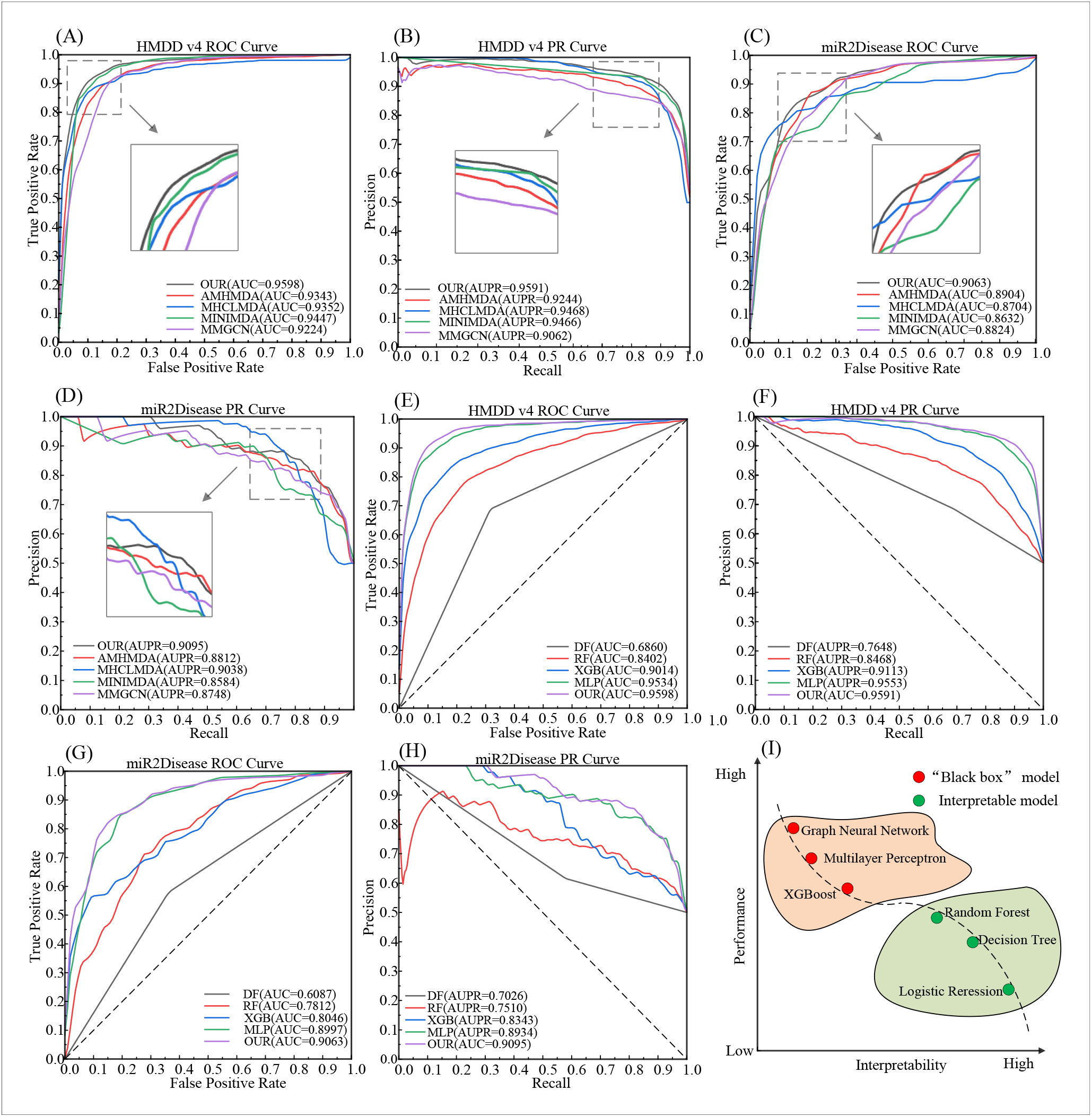
Comparison of ROC and PR curve and performance-interpretability trade-off. (A)-(D) are comparisons of iDRKAN and baseline methods on HMDD v4.0 and miR2Disease datasets. (E)-(H) are comparisons of iDRKAN and different classifiers on HMDD v4.0 and miR2Disease datasets. (I) describes the trade-off between model performance and interpretability.

#### 3.2.2 Validation of the Balance Between KAN Performance and Interpretability

With the rapid development of artificial intelligence technologies, traditional machine learning algorithms have gradually shown limitations in performance. Therefore, deep learning represented by neural networks has gained significant attention. However, the “black box” nature of deep learning models reduces the transparency of the model decision-making process. As illustrated in Fig. 4(I), from logistic regression to graph neural network, the model performance increases with the increase of complexity; However, interpretability tends to decrease. To address this, we innovatively introduce KAN as a classifier and conduct comparative experiments with classical models such as Decision Tree (DF) [40], Random Forest (RF) [41], XGBoost (XGB) [42], and Multi-Layer Perceptron (MLP) [43]. This experiment aims to verify that KAN strikes a good balance between interpretability and performance. To ensure a fair comparison, all models are based on the iDRKAN framework, with only the classifier component being different. Fig. 4(E-F) and Fig. 4(G-H) present the ROC and PR curves of iDRKAN and its different classifiers on the HMDD v4.0 and miR2Disease datasets respectively. The results indicate an increasing trend of AUC and AUPR from DF to iDRKAN on both the HMDD v4.0 dataset and miR2Disease dataset. Furthermore, Fig. 5(A-C) and Fig. 5(D-F) provide a further comparison of iDRKAN with other classifiers on HMDD v4.0 and miR2Disease datasets for other metrics. The results show a decreasing trend in ACCURACY, MCC, and F1-SCORE from iDRKAN to DF. In summary, the experimental results fully validate that iDRKAN significantly improves the prediction performance while maintaining interpretability, providing a new research perspective for MDA prediction.

**Fig. 5.**
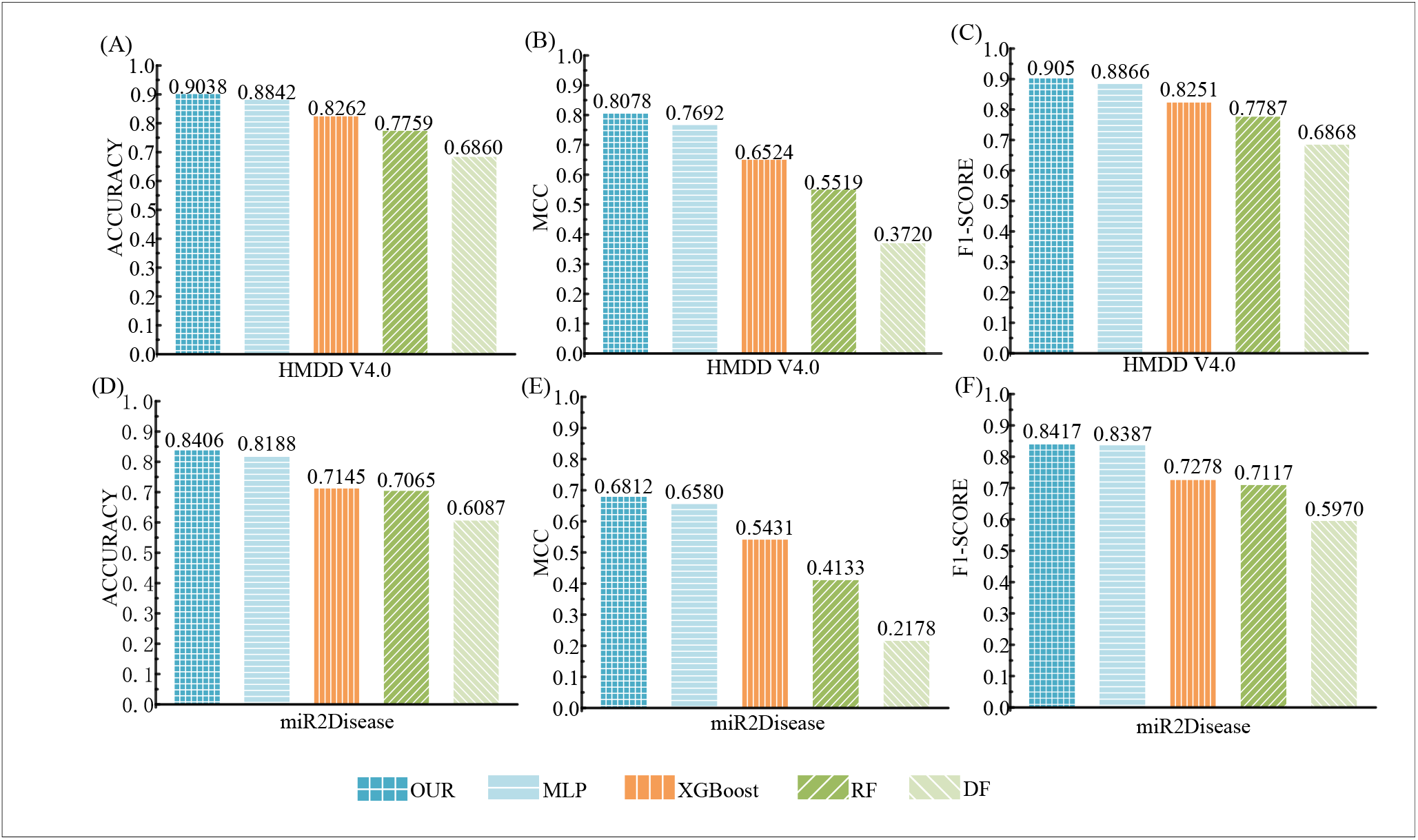
Comparison of iDRKAN and other classifiers on ACCURACY, MCC, and F1-SCORE.

### 3.3 Ablation Experiment

To evaluate the contribution of each component within the iDRKAN framework to the prediction accuracy, we construct the following four variant models: iDRKAN META (removing meta-path view), iDRKAN MCA (without multi-channel attention mechanism), iDRKAN SLA (removing semantic-layer attention mechanism), and iDRKAN CL (deleting contrastive learning strategy). The experiments are conducted on the HMDD v4.0 dataset, and the results are presented in Fig. 6. iDRKAN achieves excellent performance across metrics including AUC (0.9598), AUPR (0.9591), ACCURACY (0.9038), MCC (0.8078), and F1-score (0.9050). In contrast, iDRKAN META has the most significant decrease in AUC (0.9265), AUPR (0.9212), ACCURACY (0.8389), MCC (0.7051), and F1-score (0.8556), highlighting the importance of the deeper semantic information contained in the meta-path view. iDRKAN CL shows the second most significant performance drop, indicating the crucial role of the contrastive learning strategy in optimizing the consistency of representations across views. In summary, the ablation experiments fully confirm the positive effects of each module within iDRKAN on improving the model prediction performance.

**Fig. 6.**
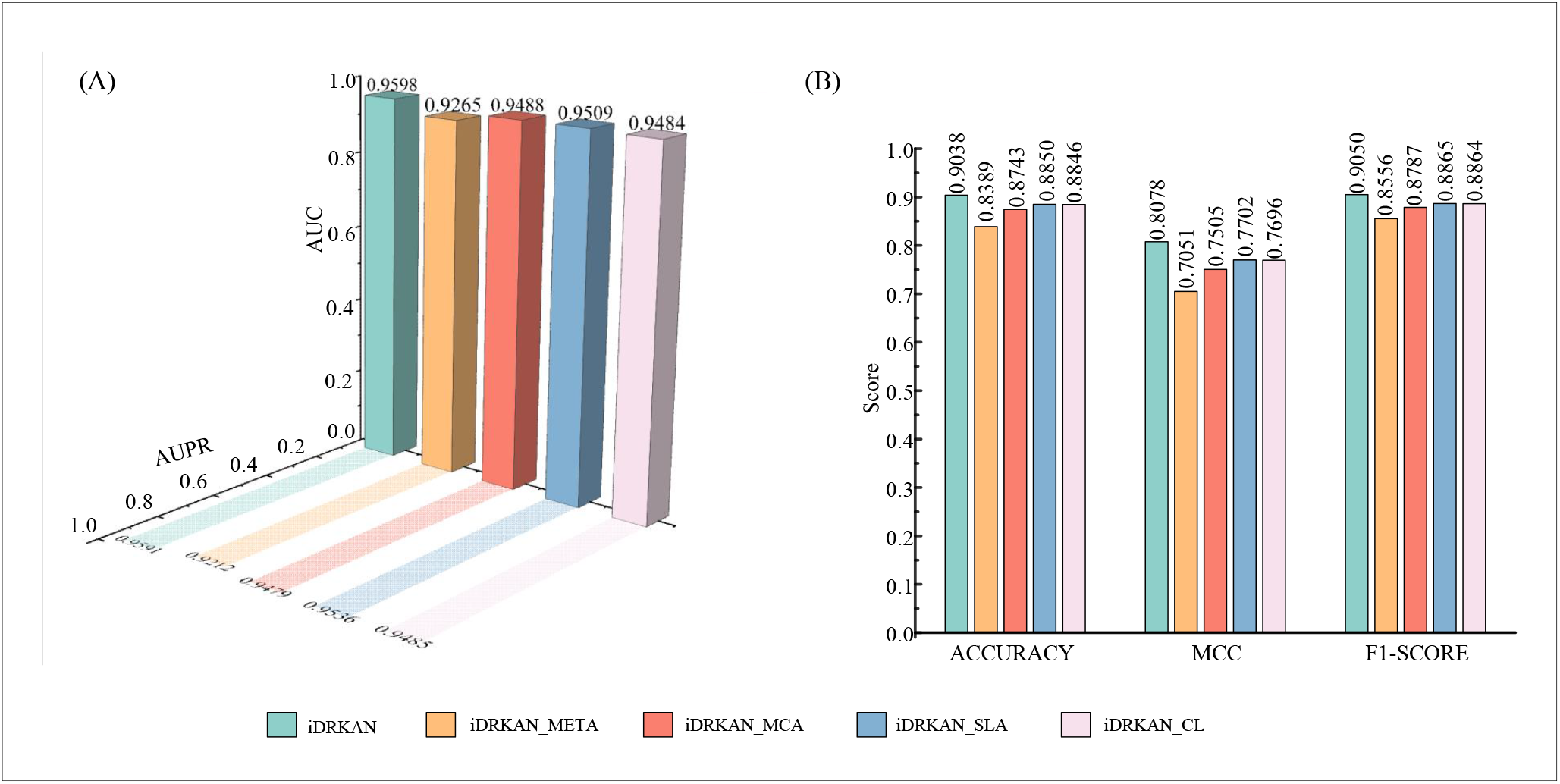
Performance comparison of iDRKAN and its variant models on the HMDD v4.0 dataset.

### 3.4 Visualization of iDRKAN Learning MDA Pair Embedding

To further validate the ability of iDRKAN to learn effective miRNA-disease pair embeddings, we perform visualization analysis on the HMDD v4.0 dataset. Specifically, we generate embedding vectors for the association pairs by applying the Hadamard product to the final miRNA and disease embeddings. According to the association existence, if there is an association between a miRNA and a disease, the pair is labeled as a positive sample; otherwise, it is labeled as a negative sample. Subsequently, the embeddings of all MDA pairs are projected into the two-dimensional space using the T-SNE method, and the results are visualized, as shown in Fig. 7. The experimental results indicate that at epoch=1, the embedding of positive and negative sample pairs is relatively chaotic. With the increase of epoch, the embedding vectors of positive and negative sample pairs are gradually clear in the two-dimensional space. Ultimately, at epoch=110, the embeddings of positive and negative sample pairs show a clear distinction. It should be noted that some regions still contain mixed red and blue points, indicating that the model still faces some challenges near the decision boundary in these regions. The above observations further confirm that iDRKAN can effectively learn discriminative MDA embeddings, thereby improving the accuracy of the model for MDA prediction.

**Fig. 7.**
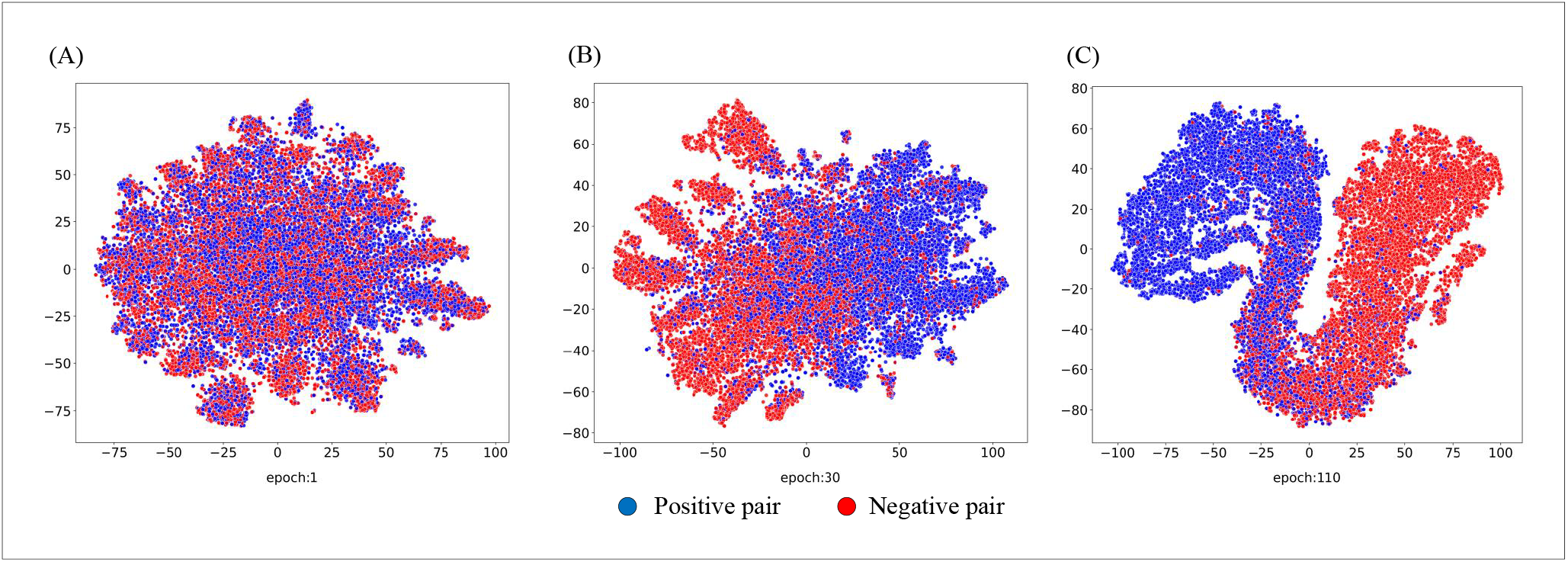
Visualization of miRNA-disease embeddings learned by GMIMDA at different epochs.

### 3.5 Case Study

To further evaluate the ability of iDRKAN to predict miRNAs associated with potential diseases, case studies are conducted on three cancers: lung, breast, and colorectal. First, the model is trained using all known associations in the HMDD v4.0 dataset to generate an association prediction score matrix. Subsequently, we screen the top 30 candidate miRNAs according to the three cancers. Finally, these candidate miRNAs are validated against the dbDEMC [44] and miRCancer [45] databases.

Lung cancer is a malignant tumor that originates in the lung tissue and is one of the most common and deadly cancers worldwide. Breast cancer is a malignant tumor occurring in breast tissue and is one of the most common cancers among women worldwide. Colorectal cancer is a malignant tumor that occurs in the colon or rectum and is a common type of cancer in the digestive tract. The validation results are shown in Tables 3 to 5. The results indicate that 30 candidate miRNAs associated with lung and colorectal cancers and 29 miRNAs associated with breast cancer are validated by dbDEMC or miRCancer database. The above three cancer case studies demonstrate that iDRKAN exhibits good performance in identifying potential MDAs.

**Table 3.** Top 30 candidate miRNAs associated with lung cancer (db1 and db2 denote dbDEMC and miRCancer))

| Rank | miRNA name | Evidence | Rank | miRNA name | Evidence |
| --- | --- | --- | --- | --- | --- |
| 1 | hsa-mir-26a | db1, db2 | 16 | hsa-mir-506 | db1, db2 |
| 2 | hsa-let-7f | db1, db2 | 17 | hsa-mir-1290 | db1, db2 |
| 3 | hsa-mir-222 | db1 | 18 | hsa-mir-181d | db1 |
| 4 | hsa-mir-335 | db1 | 19 | hsa-mir-744 | db1 |
| 5 | hsa-mir-186 | db1, db2 | 20 | hsa-mir-454 | db1 |
| 6 | hsa-mir-15b | db1, db2 | 21 | hsa-mir-376a | db1 |
| 7 | hsa-mir-212 | db1 | 22 | hsa-mir-484 | db1 |
| 8 | hsa-mir-184 | db1 | 23 | hsa-mir-433 | db1 |
| 9 | hsa-let-7e | db1, db2 | 24 | hsa-mir-888 | db1 |
| 10 | hsa-mir-363 | db1 | 25 | hsa-mir-452 | db1 |
| 11 | hsa-mir-134 | db1 | 26 | hsa-mir-762 | db1 |
| 12 | hsa-mir-346 | db1 | 27 | hsa-mir-874 | db1 |
| 13 | hsa-mir-32 | db1, db2 | 28 | hsa-mir-376c | db1 |
| 14 | hsa-mir-384 | db1 | 29 | hsa-mir-208a | db1, db2 |
| 15 | hsa-mir-421 | db1 | 30 | hsa-mir-612 | db1 |

**Table 4.** Top 30 candidate miRNAs associated with breast cancer (db1 and db2 denote dbDEMC and miRCancer)

| Rank | miRNA name | Evidence | Rank | miRNA name | Evidence |
| --- | --- | --- | --- | --- | --- |
| 1 | hsa-let-7g | db1, db2 | 16 | hsa-mir-935 | db1 |
| 2 | hsa-mir-320d | db1 | 17 | hsa-mir-587 | db1 |
| 3 | hsa-mir-346 | db1, db2 | 18 | hsa-mir-519a | db1 |
| 4 | hsa-mir-608 | db1 | 19 | hsa-mir-939 | db1 |
| 5 | hsa-mir-744 | db1 | 20 | hsa-mir-761 | db1 |
| 6 | hsa-mir-545 | db1 | 21 | hsa-mir-936 | db1 |
| 7 | hsa-mir-762 | db1, db2 | 22 | hsa-mir-596 | db1 |
| 8 | hsa-mir-888 | db1 | 23 | hsa-mir-527 | unconfirmed |
| 9 | hsa-mir-3666 | db1 | 24 | hsa-mir-588 | db1, db2 |
| 10 | hsa-mir-663 | db1 | 25 | hsa-mir-466 | db1 |
| 11 | hsa-mir-647 | db1 | 26 | hsa-mir-648 | db1 |
| 12 | hsa-mir-496 | db1 | 27 | hsa-mir-4262 | db1 |
| 13 | hsa-mir-504 | db1 | 28 | hsa-mir-4286 | db1 |
| 14 | hsa-mir-516b | db1 | 29 | hsa-mir-3180 | db1 |
| 15 | hsa-mir-636 | db1 | 30 | hsa-mir-656 | db1 |

**Table 5.**
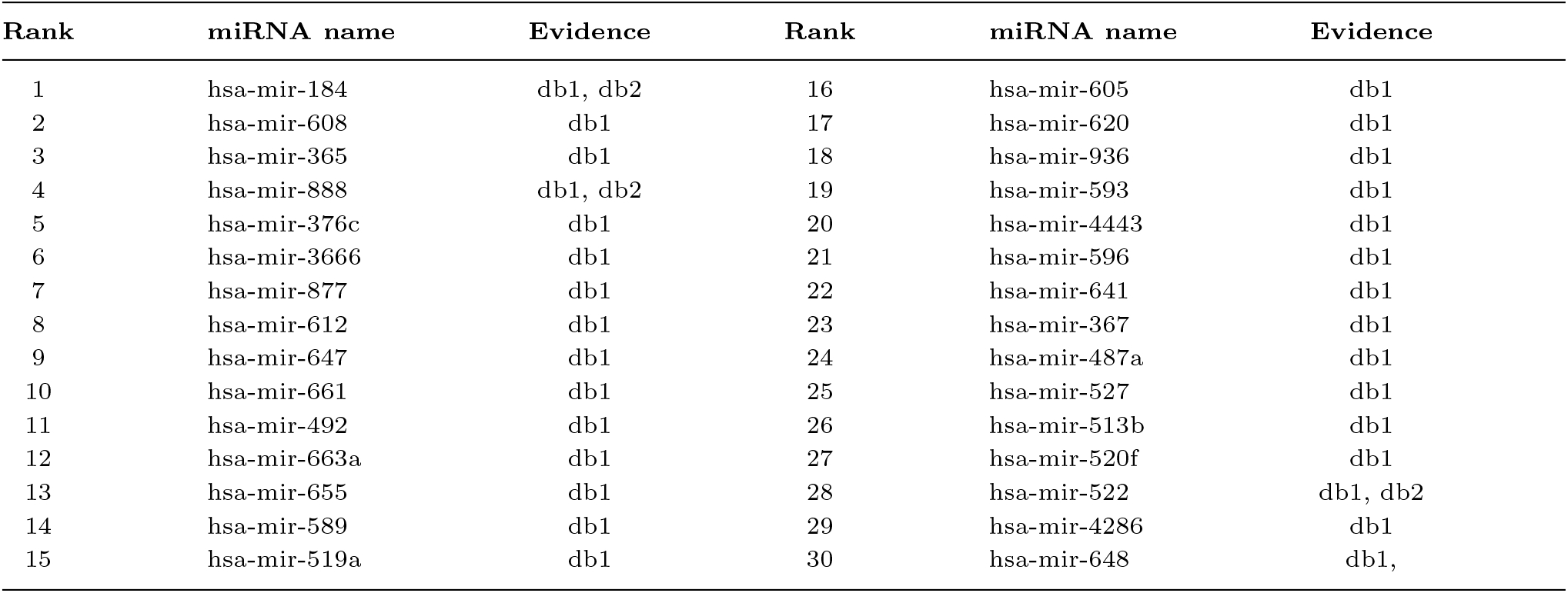
Top 30 candidate miRNAs associated with colorectal cancer (db1 and db2 denote dbDEMC and miRCancer)

| Rank | miRNA name | Evidence | Rank | miRNA name | Evidence |
| --- | --- | --- | --- | --- | --- |
| 1 | hsa-mir-184 | db1, db2 | 16 | hsa-mir-605 | db1 |
| 2 | hsa-mir-608 | db1 | 17 | hsa-mir-620 | db1 |
| 3 | hsa-mir-365 | db1 | 18 | hsa-mir-936 | db1 |
| 4 | hsa-mir-888 | db1, db2 | 19 | hsa-mir-593 | db1 |
| 5 | hsa-mir-376c | db1 | 20 | hsa-mir-4443 | db1 |
| 6 | hsa-mir-3666 | db1 | 21 | hsa-mir-596 | db1 |
| 7 | hsa-mir-877 | db1 | 22 | hsa-mir-641 | db1 |
| 8 | hsa-mir-612 | db1 | 23 | hsa-mir-367 | db1 |
| 9 | hsa-mir-647 | db1 | 24 | hsa-mir-487a | db1 |
| 10 | hsa-mir-661 | db1 | 25 | hsa-mir-527 | db1 |
| 11 | hsa-mir-492 | db1 | 26 | hsa-mir-513b | db1 |
| 12 | hsa-mir-663a | db1 | 27 | hsa-mir-520f | db1 |
| 13 | hsa-mir-655 | db1 | 28 | hsa-mir-522 | db1, db2 |
| 14 | hsa-mir-589 | db1 | 29 | hsa-mir-4286 | db1 |
| 15 | hsa-mir-519a | db1 | 30 | hsa-mir-648 | db1, |

## Conclusion

Accurately identifying disease-associated miRNAs is crucial for discovering novel biomarkers and advancing the development of targeted therapeutic drugs. With the development of intelligent computing, applying computational methods in bioinformatics has become a promising alternative. However, most of the existing methods mainly utilize contextual information in homogeneous node similarity while ignoring the complex semantic information among heterogeneous nodes. Additionally, the inherent “black-box” nature of many deep learning models further reduces the transparency of the model’s decision-making process. Therefore, we propose an interpretable computational method (iDRKAN) based on dual-graph representation learning and Kolmogorov-Arnold Network to predict MDA. iDRKAN first constructs similarity views and meta-path views, and extracts higher-order representations via GCN. Next, the context information of different similarity views is fused using MCA, while the deep semantic information in each meta-path view is integrated using SLA. Then the consistency of the representations in the two types of views is optimized based on the contrastive learning strategy. Finally, an interpretable MDA prediction is performed using the KAN with symbolic activation functions. Experimental results on two datasets show that iDRKAN outperforms the baseline approach. Case studies also validate the potential of iDRKAN in discovering potential MDA.

Although the good result of our model, there are some limitations. For example, the constructed metapath views are based on algebraic matrix operations and lack corresponding biological knowledge. Additionally, KAN relies on complex nonlinear functions (such as spline functions) to replace the fixed activation functions in traditional neural networks, which significantly increases the computational cost, especially when dealing with large-scale heterogeneous graph data. In the future, we will consider incorporating information such as genes, lncRNAs, and circRNAs into the meta-paths to enhance biological knowledge. Furthermore, MDA prediction is inherently an interaction problem, and how to model the interaction between entities to update their information is also an issue worth considering.

### Key Points

- Adding different semantic information based on meta-path views.
- Optimizing the consistency of similarity view and meta-path view representation based on contrastive learning strategy.
- Use multi-channel attention to fuse the contextual information of different similarity views; use semantic layer attention to integrate the semantic information contained in each meta-path view.
- Interpretable prediction based on Kolmogorov–Arnold Network.

## Supporting information

Similarity calculation

## Acknowledgments

The authors thank the anonymous reviewers for their valuable suggestions.

## Data Availability

https://github.com/yangfengzhuguet/iDRKAN

