## Supplementary material for "iDRKAN: Interpretable miRNA-Disease Association Prediction Based on Dual-Graph Representation Learning and Kolmogorov–Arnold Network": Similarity calculation

**Cosine similarity**

Assuming that the miRNA-disease association matrix is denoted as
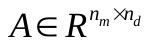
 , where
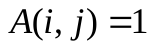
if there is an association between the miRNA
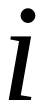
 and the disease
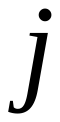
 , and
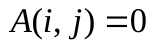
 otherwise. Here,
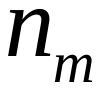
 and
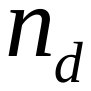
 represent the number of miRNAs and diseases, respectively. The cosine similarity
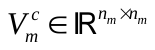
 between the miRNA
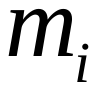
 and
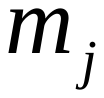
 is calculated as follows:

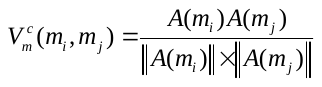

where
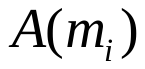
 and
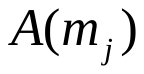
 represent the
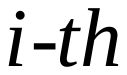
 and
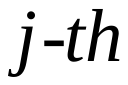
 rows of the miRNA-disease association matrix, respectively.

Similarly, the cosine similarity of diseases is denoted as
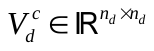
 .

**Semantic similarity of diseases**

We downloaded the latest MeSH from the U.S. National Library of Medicine to aid in the calculation of semantic similarity between diseases [1]. First, we standardized the disease names in the benchmark dataset to ensure consistency with MeSH terminology. Subsequently, we constructed a directed acyclic graph (DAG) to compute the semantic similarity between different diseases. In this DAG, nodes represent individual diseases, while edges depict their relationships [2]. Here, we define two types of semantic similarity for disease
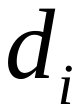
 and
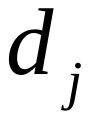
：
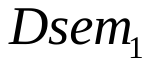
 and
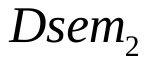
 .

The directed acyclic graph of disease
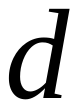
 is denoted as
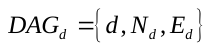
, where
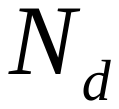
 represents the disease
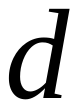
 and its set of neighboring nodes and
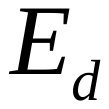
 denotes the associations among different diseases. If a disease
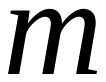
 exists in
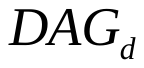
 , the semantic contribution of
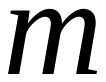
 to
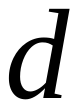

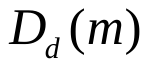
 is defined as follows:

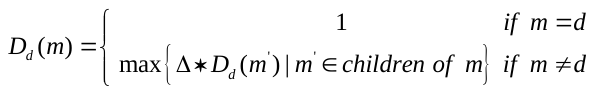

where
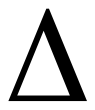
 represents the semantic contribution factor and the value is set to 0.5 according to a previous study [3].

Next, the semantic value for each disease is calculated based on the semantic contributions from all neighboring nodes. For a given disease
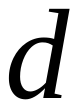
 , the semantic value
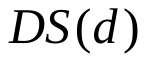
 is determined as follows:

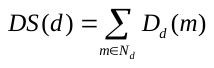

In the DAG, if two diseases overlap more, the more ancestors they share, indicating that they have higher semantic similarity. Consequently, the semantic similarity
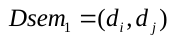
 between diseases
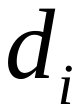
 and
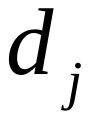
 is defined as follows:

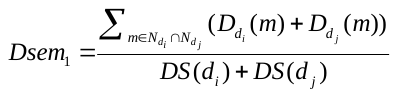

Based on hierarchical relationships,
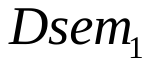
 primarily quantifies diseases according to their relative position within the DAG. However, this approach does not consider quantitative differences among diseases. Considering that rare diseases often exert a greater impact than common diseases, the following formula is employed to quantify the contribution of disease
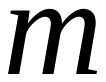
 to disease
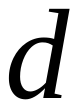
 :

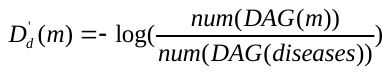

where
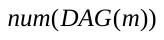
 represents the number of diseases
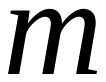
 in the DAG, while
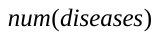
 indicates the total number of diseases. Accordingly, another semantic similarity,
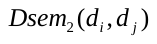
 , between disease
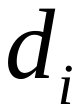
 and
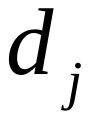
 is computed as:

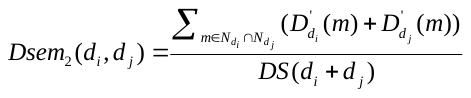

Finally,
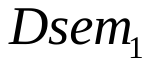
 and
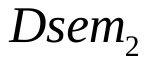
 are combined to obtain the final disease semantic similarity
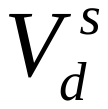
 :

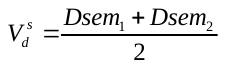

**GIP kernel similarity**

To evaluate the topological similarity between network nodes, the GIP kernel similarity matrix was constructed between miRNAs and diseases using the miRNA-disease association matrix [4]. For example, consider the GIP similarity
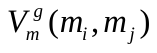
 between miRNA
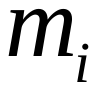
 and
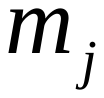
. First,
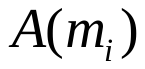
 is defined as the disease binary vector associated with miRNA
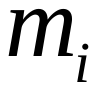
 , which corresponds to the
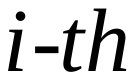
 row of the miRNA-disease association matrix. Similarly,
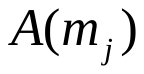
 represents the
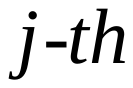
 row of the miRNA-disease association matrix. Thus, the GIP similarity
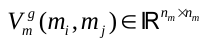
, calculated using a Gaussian kernel function, is defined as follows:

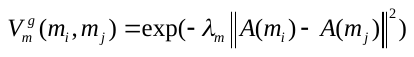

where
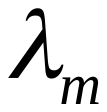
 denotes the bandwidth of the kernel, which is calculated as:

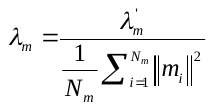

where
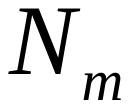
 denotes the total number of miRNAs, and
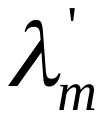
 represents the normalization constant, which is set to 1 according to the previous study [5].

Similarly, the GIP similarity for diseases,
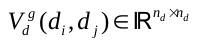
, can be obtained.

**miRNA functional similarity**

According to the study by Wang et al. [2], miRNAs with similar functions are often associated with similar diseases. We calculated the functional similarity of miRNAs based on known miRNA-disease associations and disease similarities. For miRNA
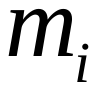
 and
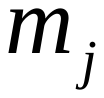
,
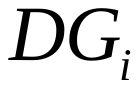
 and
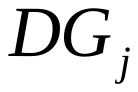
 are defined as the set of diseases associated with
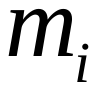
 and
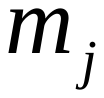
 based on the known miRNA-disease associations respectively.

To quantify the similarity between two disease groups, the similarity between a single disease and a disease group must first be calculated. Given a disease group
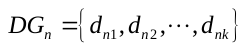
, the similarity between a single disease
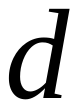
 and the group
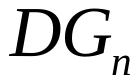
 is defined as follows:

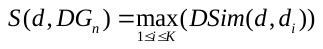

where
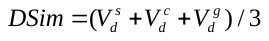
 represents the combined similarity matrix for the two diseases, and
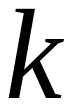
 denotes the number of diseases in
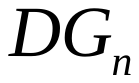
 .

Consequently, the functional similarity
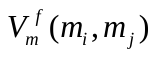
 between miRNA
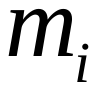
 and
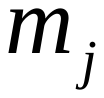
 is determined as follows:

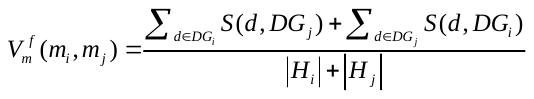

where
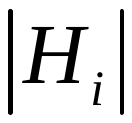
 and
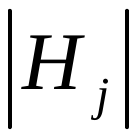
 denote the number of diseases in
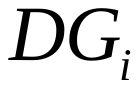
 and
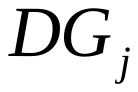
 respectively.
